# Transcriptional response of individual Hawaiian *Culex quinquefasciatus* mosquitoes to the avian malaria parasite *Plasmodium relictum*

**DOI:** 10.1101/2022.02.10.479890

**Authors:** Francisco C. Ferreira, Elin Videvall, Christa M. Seidl, Nicole E. Wagner, A. Marm Kilpatrick, Robert C. Fleischer, Dina M. Fonseca

## Abstract

*Culex quinquefasciatus*, the mosquito vector of avian malaria in Hawai□i, became established in the islands in the 1820s and the deadly effects of malaria on endemic bird species have been documented for many decades. To evaluate the gene expression response of the mosquito to the parasite, we let the offspring of wild-collected Hawaiian *Cx. quinquefasciatus* feed on a domestic canary infected with *Plasmodium relictum* GRW4 freshly isolated from a wild-caught Hawaiian honeycreeper. Control mosquitoes were fed on an uninfected canary. We sequenced the individual transcriptomes of five infected and three uninfected individual mosquitoes at three different stages of the parasite life cycle: 24 h post feeding (hpf) during ookinete invasion; 5 days post feeding (dpf) when oocysts are developing; 10 dpf when sporozoites are released and invade the salivary glands. Differential gene expression analyses showed that during ookinete invasion (24 hpf), genes related to oxidoreductase activity and galactose catabolism had lower expression levels in infected mosquitoes compared to controls. Oocyst development (5 dpf) was associated with reduced expression of a gene with a predicted innate immune function. At 10 dpf, infected mosquitoes had reduced expression levels of a serine protease inhibitor. Overall, the gene expression response of Hawaiian *Culex* exposed to a *Plasmodium* infection intensity that occur naturally in Hawaii was low, but more pronounced during ookinete invasion. The low fitness costs often documented in *Culex* infected with avian *Plasmodium* likely reflect the relatively small transcriptional changes observed in mosquito genes related to immune response and nutrient metabolism.

## Introduction

Avian malaria in Hawai□i, a mosquito-borne disease caused by *Plasmodium relictum*, is an emblematic example of the deadly impacts that an invasive pathogen can pose to wildlife (van Riper et al., 1986; Warner, 1968). While the abundance and distribution of endemic Hawaiian honeycreepers (Fringillidae: Drepanidinae) have been dramatically impacted by habitat modification and the introduction of invasive vertebrates (Pratt et al. 2009), avian malaria is one of the main factors that drove several honeycreeper species to extinction or endangerment (van Riper et al., 1986; Warner, 1968). Avian malaria in Hawai□i is transmitted by the southern house mosquito (*Culex quinquefasciatus*), a highly competent vector (LaPointe et al., 2005, 2010) that was introduced first to Lahaina, Maui presumably in 1826 (Dine, 1904). It is now clear, however, that multiple *Cx. quinquefasciatus* introductions have occurred in Hawai□i (Fonseca et al., 2000) and the initial American-derived mosquitoes were later replaced by populations originating in the southwest Pacific region (Aardema et al., 2021; Fonseca et al., 2006). Although *Cx. quinquefasciatus* had been widespread across Hawai□i since the late 1800s, it was not until the 1950s that malaria was confirmed as a cause of mortality for native birds (Warner, 1968).

Low- and mid-elevation areas across Hawai□i provide habitat, temperature, and humidity favorable for both mosquito proliferation and *Plasmodium* development within the vector, effectively limiting honeycreeper populations mostly to high elevation habitats (Atkinson & LaPointe, 2009). However, some populations of Hawai□i ‘Amakihi (*Chlorodrepanis virens*) have persisted in low-elevation areas (Woodworth et al., 2005), and survive infections (Atkinson et al., 2013; Cassin-Sackett et al., 2019). Most of the birds in these low-elevation native bird populations are susceptible to *Plasmodium* and become infectious to mosquitoes (McClure et al., 2020; Woodworth et al., 2005). As a consequence, higher mosquito infection rates are observed in areas with higher densities of native birds, suggesting that the presence of these species increases malaria transmission, which in turn may negatively affect the survival of other more susceptible native Hawaiian bird species in low elevation areas (McClure et al., 2020). Reducing malaria transmission via suppression of mosquito populations and/or a reduction of vector competence of local *Cx. quinquefasciatus* could contribute to the reestablishment of stable Hawaiian bird populations in lowland areas, but mosquito control methods have been unsuccessful at protecting Hawaiian honeycreepers from malaria, partly because of difficult access to mosquito breeding sites and the potential of non-target effects of insecticides (Pratt 2009). Therefore, other methods to reduce or block *Plasmodium* transmission are needed (Paxton et al., 2018). One proposed strategy is the development of a synthetic gene drive system to make mosquitoes refractory or resistant to the parasite (Nishimoto, 2019), and recent developments of CRISPR tools for *Cx. quinquefasciatus* (Feng et al., 2021) have opened exciting new opportunities for genetically engineered mosquitoes to be potentially released as a conservation option in the future. This approach, however, requires information about mosquito genes directly related to *P. relictum* invasion, development, and transmission.

*Plasmodium* parasites undergo a complex life cycle within their vectors, and the main developmental stages are relatively conserved among species that infect birds and mammals (Ferreira et al., 2020). Blood feeding female mosquitoes ingest gametocytes, and these parasites develop to gametes that form a motile ookinete as early as 16 h post mosquito feeding (van Riper et al., 1986). Ookinetes cross the blood meal peritrophic membrane and attach to the mosquito midgut epithelium, initiating the interaction between the parasite and its potential vector. Ookinetes then actively cross the midgut epithelia and continue their development. This migration is a critical bottleneck for *Plasmodium* development because an infected mosquito may activate immune-related genes to kill ookinetes (Rhodes & Michel, 2017). Ookinetes that survive develop into oocysts where sporozoites are produced via sporogony using mosquito resources such as lipids, carbohydrates, and amino acids (Shaw et al., 2021). Mature oocysts burst and release thousands of sporozoites into the mosquito hemocoel, a subset of which reaches the salivary glands and cross its epithelial layer, where they remain until the mosquito takes another blood meal. During the probing process on the host, sporozoites in the mosquito saliva are injected into the skin where their journey continue in the vertebrate host (Valkiūnas & Iezhova, 2017).

Recent studies conducted on Hawaiian honeycreepers unveiled bird genes linked to immune functions that may be under selection in ‘amakihi populations that developed higher survival to malaria (Cassin-Sackett et al., 2019) and explored the transcriptome of *P. relictum* blood stages in experimentally infected ‘amakihis (Videvall et al., 2021). However, the genetic basis of the response of vectors to *Plasmodium* parasites in Hawai□i or in other geographical areas is still unknown, despite avian malaria being globally widespread (Fecchio et al., 2021). Acquiring this information will allow the comparison of the responses of vectors of mammalian (*Anopheles* mosquitoes) and avian (*Culex* mosquitoes) *Plasmodium* parasites. These comparisons are particularly interesting given that the subfamilies Anophelinae and Culicinae diverged around 200 million years ago (Reidenbach et al., 2009), while the split between avian and primate/rodent *Plasmodium* is estimated to have occurred around 45 million years ago (Pacheco et al., 2018). Using avian malaria parasites and four mosquito genera as models, Huff (1927) suggested that the vector’s immune response plays a key role in *Plasmodium* refractoriness. While his study forms the basis to our understanding of vector competence to malaria, the vector’s responses to *Plasmodium* have so far been studied mainly in human and rodent malaria systems using *Anopheles* mosquitoes.

We analyzed the transcriptional response of representative Hawaiian *Cx. quinquefasciatus* exposed to Hawaiian *P. relictum*. Mosquitoes were divided into two groups: one that fed on a *P. relictum*-infected canary (*Serinus canaria*), and another group that fed on an uninfected canary to serve as a baseline for analyses. *Plasmodium* parasites induce gene expression changes in *Anopheles* mosquitoes during ookinete invasion, oocyst development, and when sporozoites spread into the vector hemolymph and invade the salivary glands of the vector (Rhodes & Michel, 2017). Therefore, we sequenced and quantified the transcriptome of single *Cx. quinquefasciatus* to address individual response to *P. relictum* at these three critical time points for *Plasmodium* development within the vector.

## Methods

### *Plasmodium relictum* isolation and *Culex quinquefasciatus* collection

We isolated *Plasmodium relictum* (lineage GRW4) from a single Hawai□i ‘amakihi (*Chlorodrepanis virens*) captured in Nanawale Forest Reserve (19°32’14.4”N 154°54’11.6”W) in February 2020. This is a low elevation area (98 meters above sea level) where Hawai□i ‘amakihi constitute 20% of the avian community (McClure et al., 2020).

Around 100 µl of blood was collected from the brachial vein into a 1 ml syringe containing 14 µl of Citrate-phosphate-dextrose solution with adenine (Sigma-Aldrich, St. Louis, MO, USA), as described by Carlson et al. (2016). The sample was stored at 4ºC for 48 h before being inoculated into the pectoral muscle of an uninfected domestic canary, *Serinus canaria*. Ten days post inoculation, this canary developed a parasitemia of 1.72% and we collected 100 µl of blood as described above. That blood sample was shipped on ice from Hawai□i to Rutgers University in New Jersey and was intraperitoneally inoculated into two *Plasmodium*-free canaries 36 h after it had been collected.

To obtain the experimental mosquitoes, we collected 30 mosquito egg rafts in Captain Cook, Hawai□i Island (19°27’40.6”N 155°53’47.4”W – 4.5 km from sampling site 17, Kealakekua, described in Fonseca et al. 2006) at 204 meters above sea level. The egg rafts were shipped to Rutgers University and maintained in an incubator at 26 ºC, with 70-80% relative air humidity under a photoperiod regime of 13L:11D. After hatching, larvae were fed with commercial ground fish food until pupation. Pupae were transferred to mosquito cages kept in the same incubator, and emerging adults were maintained on a 10% sucrose solution. Female mosquitoes were fed on uninfected canaries to create an F1 generation of mosquitoes under laboratory conditions. For all blood feeding procedures, canaries were immobilized in a plastic cylinder, in which only their legs are accessible to mosquitoes (Kazlauskienė et al., 2013), and placed inside the mosquito cages maintained in the incubator. Experimental procedures were approved by Rutgers University Institutional Animal Care and Use Committee (PROTO201900075) and by University of California Santa Cruz IACUC (protocol kilpm2003). Permits for bird sampling include U.S. Department of the Interior Bird Banding Laboratory permit #23600, Hawai□i State Department of Land and Natural Resources Protected Wildlife Permit WL 19-23 Amend 01, Hawai□i State Access and Forest Reserve Special Use Permit. Mosquitoes and *Plasmodium* isolates were transported from Hawai□i to New Jersey under USDA-APHIS permits number 140413 and 141156.

### Mosquito infections and assessment of parasite development

In our experiments, 7-8 day-old female mosquitoes were allowed to feed for one hour on a single infected canary. This bird had been experimentally inoculated 18 days prior with a second-passage of *Plasmodium*, and was sampled every 2-3 days after the fifth day post infection. Blood samples were assessed by PCR (Hellgren et al., 2004) and light microscopy (Valkiūnas, 2005) for the presence of *Plasmodium*. Mosquitoes from the same batch were allowed to feed on an uninfected canary, also for 1 h, to serve as negative controls for baseline transcriptome data. Fully engorged mosquitoes from infected and control groups were transferred to separate cages and kept under the same conditions as described above. For transcriptome analyses, we processed mosquitoes at three time points of *P. relictum* development: (1) midgut invasion by ookinetes at 24 hours post feeding (hpf); (2) oocyst development at 5 days post feeding (dpf); and (3) sporozoite release into the hemocoel and invasion of the salivary glands at 10 dpf. Previous experiments (data not shown) conducted under the same conditions revealed that ookinetes were present in the midgut 24 hpf; immature oocysts were present in the midgut at 5 dpf, but no sporozoites were detected in the salivary glands; mosquitoes at 8 dpf harbored sporozoites in the salivary glands and were already capable of infecting domestic canaries via blood feeding. We added two extra incubation days at the last time point for the transcriptome experiments to allow for longer oocyst development and sporozoite invasion of salivary glands. At each timepoint, we collected seven to nine mosquitoes with a battery-operated aspirator and immediately transferred them to an insulated box containing dry ice. After this step, we quickly transferred individual mosquitoes to screw-cap microtubes on dry ice that were stored at -80º C until RNA extraction. Mosquito challenges were conducted in a USDA-APHIS inspected BSL-2 insectary.

### Mosquito dissection and parasite quantification

Two additional mosquitoes from the infected group were dissected at each timepoint for *Plasmodium* evaluation. At 24 hpf, we dissected their midguts onto glass slides and mixed the visible blood meal with saline solution (0.9% NaCl). These preparations were air-dried, fixed with absolute methanol, and stained with 10% Giemsa solution for 60 minutes. We dissected two mosquitoes at 5 dpf and at 10 dpf by individually pulling their midguts onto glass slides containing saline solution. A glass coverslip was gently placed over the midguts and subsequently examined with a microscope under 100 x and 400 x magnification to detect oocysts. After quantification of oocysts, we transferred each midgut to individual microtubes that were kept at -20º C. Using sterile dissecting needles, we extracted the salivary glands from these mosquitoes onto individual glass slides containing saline solution forming thin smears. Head and thorax remnants of dissected mosquitoes were transferred to individual microtubes and stored at -20º C. Slides with salivary glands preparations were air-dried, fixed with absolute methanol, and stained with 4% Giemsa solution for 60 minutes. We used PCR targeting the *Plasmodium cytb* locus (Hellgren et al., 2004) to test the midguts and thorax remnants of mosquitoes dissected at 5 dpf and 10 dpf for the presence of *Plasmodium* and we sequenced the positive samples to confirm the parasite identity.

### RNA extraction, library preparation and sequencing

We extracted RNA from individual mosquitoes using TRIzol^®^ (Invitrogen, Carlsbad, CA, USA) followed by column purification using RNeasy mini kit^®^ (QIAGEN, Hilden, Germany). First, we added 500 μL of TRIzol and one 3-mm glass bead to the microtubes with mosquitoes and then homogenized the samples in a TissueLyser^®^ (QIAGEN, Hilden, Germany) at 30 Hz for 3 min. We then added an additional 600 μL of TRIzol to the samples, and incubated them at room temperature for 3 min. After this step, we added 220 μL of chloroform, shook the tubes by hand vigorously for 15 seconds and incubated the samples at room temperature for 3 minutes. We centrifuged the samples at 12,000 g for 15 minutes in a cooled centrifuge and transferred 650 μL of the upper aqueous phase to a new microtube. We added an equal volume (650 μL) of 70% ethanol to the supernatant and transferred this mixture to Qiagen RNeasy^®^ mini columns (QIAGEN, Germantown, MD, USA). We performed downstream processes following the manufacturer’s instructions with the additional DNase I digestion step. The RNA was suspended in 50 μL of RNAse-free water and checked using a NanoDrop™ 2000 (Thermo Scientific, Waltham, MA, USA). We kept the tubes containing the precipitate material with TRIzol^®^ for DNA extraction conducted following the manufacturer’s protocol to confirm via PCR that mosquitoes had acquired infection after feeding on the infected canary.

We used 200 ng of total RNA for each library preparation. mRNA was isolated using NEBNext Poly(A) mRNA Magnetic Isolation Module (New England Biolabs, Ipswich, MA, USA) following the manufacturer’s protocol. The isolated mRNA was then used to make transcriptome libraries using the NEBNext Ultra II Directional RNA Library Prep Kit with NEBNext Multiplex Oligos for Illumina (Index Primers sets 1 - 4) according to the manufacturer’s protocol. Library quality was assessed on an Agilent 2100 Bioanalyzer using High Sensitivity DNA reagents and chips (Agilent Technologies, Santa Clara, CA, USA). Library concentration was measured with a Qubit 2.0 Fluorometer using a dsDNA BR Assay Kit (Life Technologies, Carlsbad, CA, USA). All 48 libraries were pooled in an equimolar fashion and submitted to Genewiz (South Plainfield, NJ, USA) for paired-end sequencing (2×150 bp) in one lane of Illumina HiSeq 2500.

## Data analyses

Raw reads were trimmed of adaptors and quality-filtered using Trimmomatic (ver. 0.39; Bolger et al., 2014). Read quality was assessed using FastQC (ver. 0.11.9; www.bioinformatics.babraham.ac.uk/projects/fastqc) together with MultiQC (ver. 1.9; Ewels et al., 2016). A total of 286 million 150 bp paired-end reads passed quality control, with an average of 12M reads per sample. We used HISAT2 (ver. 2.2.1; Kim et al., 2019) to map trimmed reads to the concatenated genomes of *Culex quinquefasciatus* (Johannesburg strain, ver. VectorBase 48; https://vectorbase.org/vectorbase/app), *Plasmodium relictum* (PrelictumSGS1-like/ PlasmoDB 48; https://plasmodb.org/plasmo/app/) (Böhme et al., 2018) and domestic canary (https://www.ncbi.nlm.nih.gov/assembly/GCF_007115625.1). The three genomes were concatenated to allow potential reads from mosquitoes, parasites and birds to map to their respective genomes. In a second step, we kept only the reads that mapped to the *Cx. quinquefasciatus* genome using Samtools (ver 1.10; Li et al., 2009), to ensure only the mosquito genes were analyzed. The reads were then counted using HTSeq (ver. 0.11.1; Anders et al., 2015), and differential gene expression analyses were performed with DESeq2 (ver. 1.28.1; Love et al., 2014) in R (ver. 4.0.3; R Development Core Team, 2020). Counts were normalized for library size differences using the geometric mean and modeled with a negative binomial distribution. To visualize samples on a PCA without bias we performed Variance Stabilizing Transformation of counts according to the manual. Because mosquito transcriptomes had been sequenced individually, and not pooled as in most studies, we could account for the biological variation between individuals in the analyses. Differentially expressed genes (DEG) were compared between infected and control mosquitoes within each time point using Benjamini and Hochberg false discovery rate to correct for multiple testing. Genes were considered significantly differentially expressed using the default DESeq2 threshold of p-adjusted values < 0.1.

We conducted Gene Ontology (GO) analyses of the DEGs in VectorBase 54 (Giraldo-Calderón et al., 2015) using the domains biological processes and molecular functions. Functional groups with a Benjamini-Hochberg false discovery rate (FDR) of < 0.1 were considered as statistically enriched. The list of enriched GO terms was filtered for redundant terms manually and through REVIGO (Supek et al., 2011). The final list was used for visual representations as described by Bonnot et al. (2019).

## Results

### *Plasmodium relictum* development in mosquitoes

The infected canary was *Plasmodium*-positive by both PCR and microscopy for the first time at 7 days post infection (dpi), displaying an initial parasitemia of 0.02% and reaching a peak parasitemia of 1.11% at 16 dpi. On the day of the mosquito exposure experiments (18 dpi), the bird had a parasitemia of 0.92%, with 0.12% of the total erythrocytes infected with mature gametocytes (the sexual stages that give rise to male and female gametes in the mosquito midgut).

We did not detect ookinetes in the midguts of the two mosquitoes dissected at 24 hpf. At 5 dpf, we identified one or two oocysts in the midgut of each mosquito, and no sporozoites in the salivary glands. Thorax remnants from both dissected mosquitoes at 5 dpf were PCR-negative, confirming that sporozoites were not present in the mosquitoes’ salivary glands at that time point. At 10 dpf, we identified three oocysts in the midgut preparations of both mosquitoes, as well as low densities of sporozoites in the salivary glands. Midguts from 5 dpf and 10 dpf and thorax remnants at 10 dpf were PCR-positive and *cytb* sequencing confirmed the expected GRW4 lineage identity of *Plasmodium relictum* present in the infected mosquitoes. Zero mosquitoes died during the 10-day period of the experiment.

For the mosquitoes that were prepared for RNA extraction, *Plasmodium* infection was confirmed by PCR in all samples at 24 hpf and 10 dpf. One mosquito at 5 dpf was PCR-negative and was substituted with a backup mosquito that was positive by PCR. As a result, infection was confirmed in all mosquitoes exposed to *Plasmodium* for the transcriptome analyses.

### Overall *Culex* transcriptome response to *Plasmodium relictum*

Most of the transcriptome variation in the PCA (62%) was driven by the response of 24 h post feeding mosquitoes, regardless of their *Plasmodium* infection status and there was no clear clustering of mosquitoes at 5 dpf or at 10 dpf or based on their infection status (Figure 1). Two genes were differentially expressed at more than one time point: CPIJ018704 (Transmembrane protein 104) had higher expression levels at 5 dpf and lower expression levels at 10 dpf in infected mosquitoes compared to the control group; CPIJ010933 (uncharacterized protein) had lower expression levels at 5 dpf, but higher expression levels at 10 dpf.

**Figure 1:**
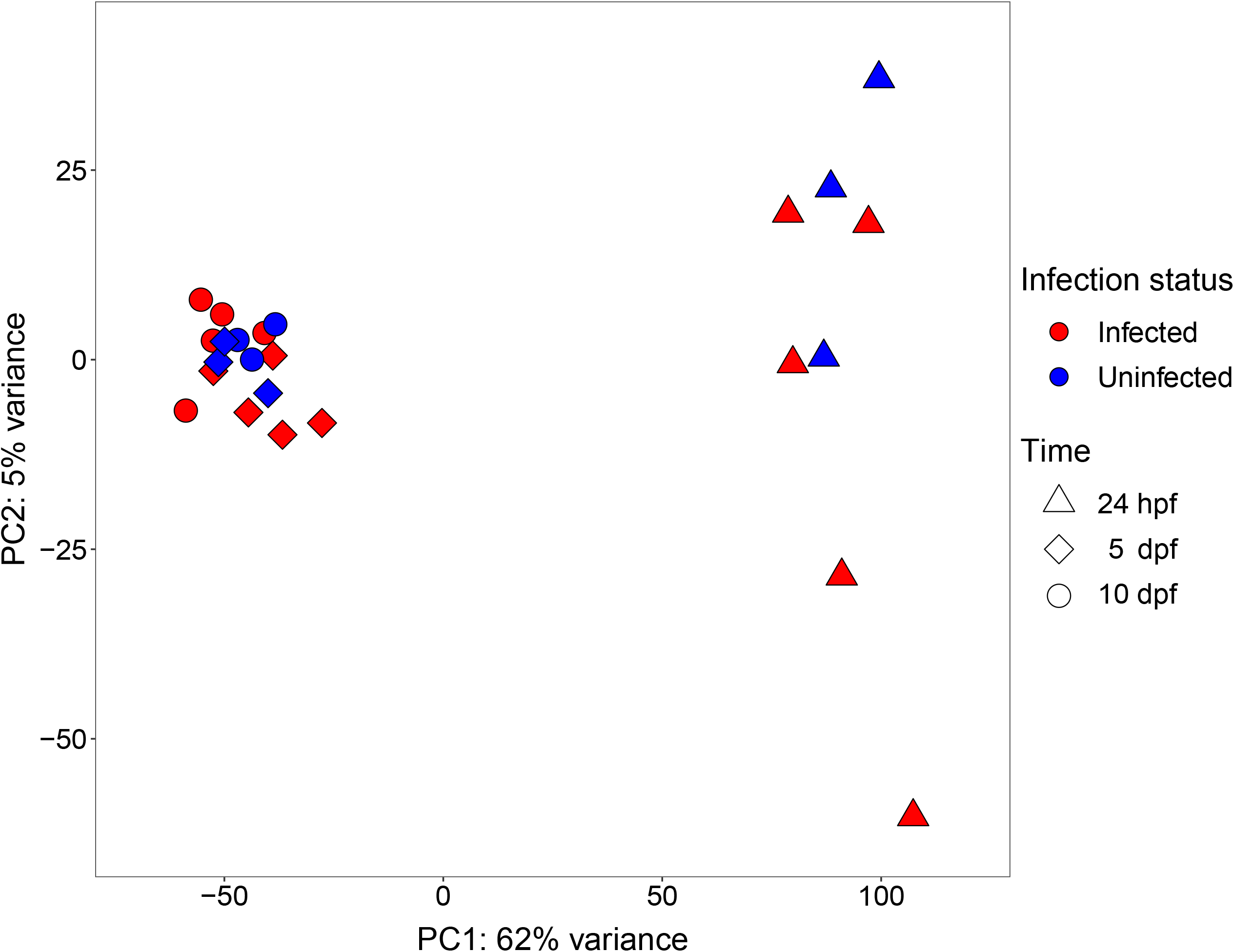
Principal component analysis (PCA) of transcriptome variation in *Culex quinquefasciatus* mosquitoes at three time point after feeding on either a negative-control canary (Uninfected) or a canary infected with a Hawaiian strain of *Plasmodium relictum* GRW4 (Infected). Time points analyzed were: 24 h post feeding (24 hpf = ookinete invasion of mosquito midgut), 5 days post feeding (5 dpf = oocyst development), and 10 days post feeding (10 dpf = oocyst maturation, sporozoite release and invasion of salivary glands). Most of the variance was explained by mosquitoes 24 hpf regardless of *Plasmodium* infection status.

### Transcriptional response at 24 h post blood feeding – ookinete invasion

During the initial *Plasmodium* invasion, 24 h after feeding on an infected bird, 109 genes were differentially expressed in the infected mosquitoes compared to the control group, with 53 genes having higher expression levels, and 56 genes having lower expression levels (Figure 2a, Table S1). Infected mosquitoes had higher expression levels of genes related to cytoskeleton organization (CPIJ001347, CPIJ003238 and CPIJ013059) and of a Down syndrome cell adhesion molecule gene paralog (CPIJ007041). Gene ontology analyses using genes with lower expression levels revealed enriched biological processes that include galactose catabolism and acetyl-CoA biosynthesis (Figure 3). Enriched molecular processes included oxidoreductase activity, serine-type endopeptidase activity and ion binding for a variety of metals. Almost significant (FDR = 0.107) enriched GO terms of genes with higher expression levels in infected mosquitoes were related to molecular functions such as calcium transportation, cobalamin binding and peptidoglycan muralytic activity (Table S2, Figure S1).

**Figure 2.**
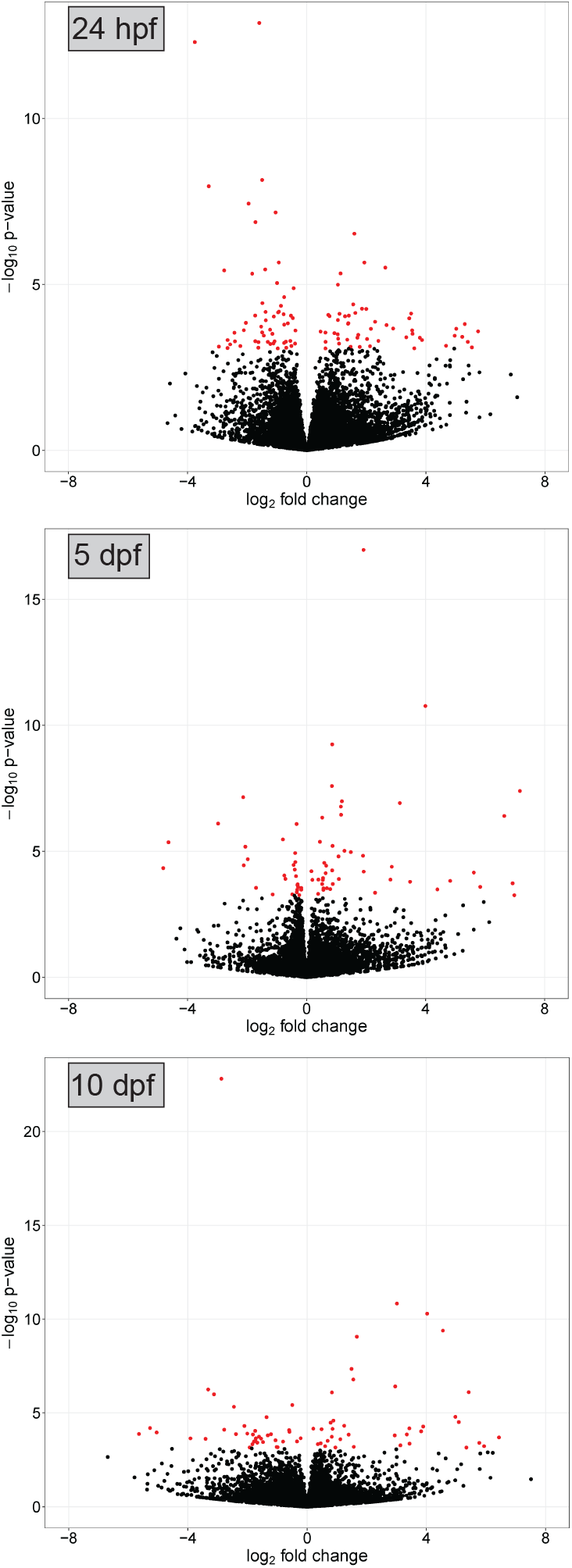
Volcano plots showing *Culex quinquefasciatus* gene expression at three time points after feeding on a canary infected with a Hawaiian strain of *Plasmodium relictum*. Mosquitoes that fed on an uninfected canary were used as negative controls, and time points analyzed are described in the legend of Figure 1. Red circles illustrate significant genes with lower (negative) or higher (positive) expression levels in infected mosquitoes compared to uninfected ones. Black circles illustrate nonsignificantly expressed genes.

**Figure 3.**
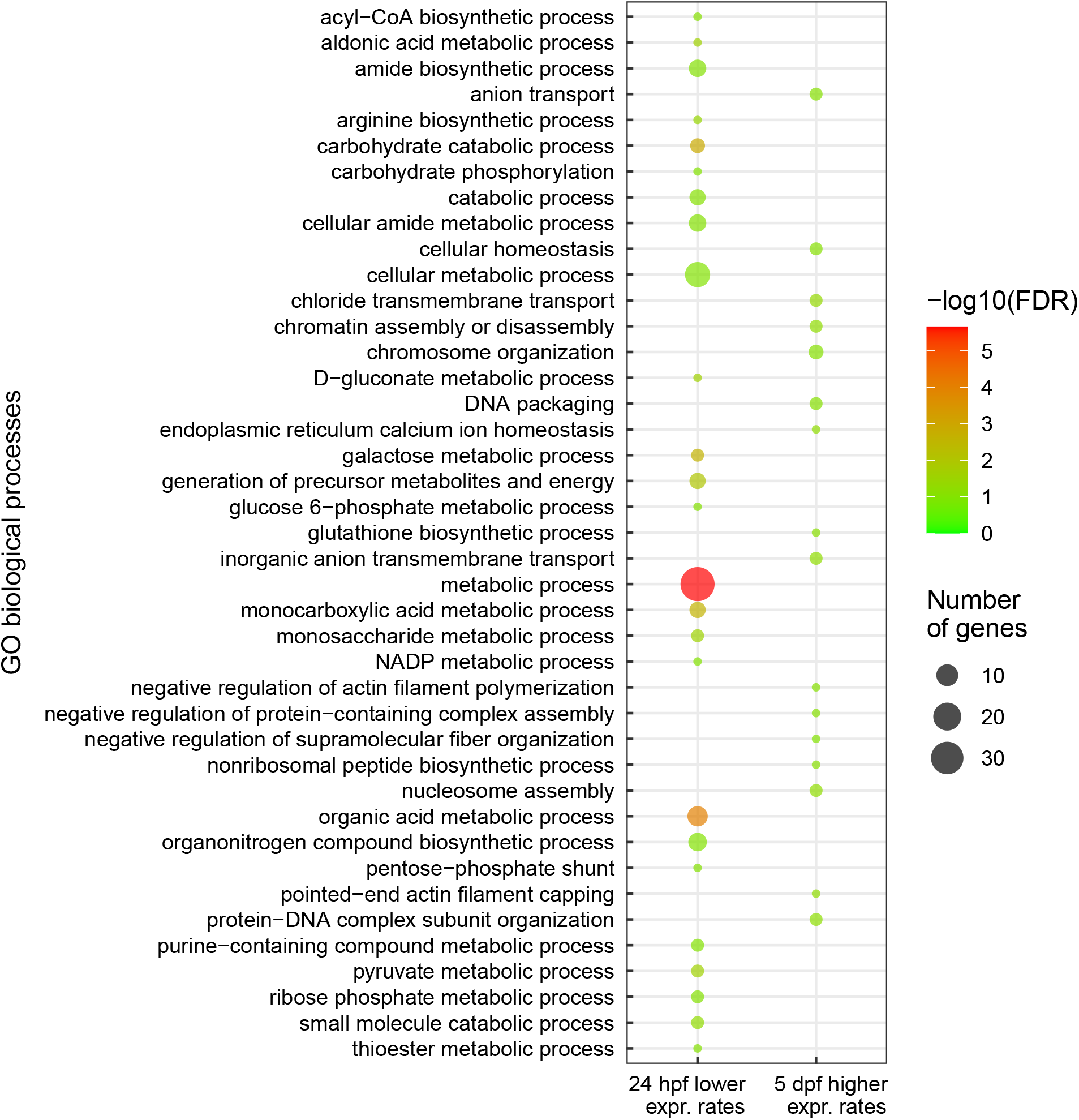
Enriched Gene Ontology terms for biological processes among differentially expressed genes in infected mosquitoes compared to uninfected ones. Only time points with statistically significant enriched GO terms are displayed. hpf = hours post feeding, dpf = days post feeding.

### Transcriptional response at 5 days post blood feeding – oocyst development

During oocyst development at 5 days dpf, before sporozoites are released into the mosquito haemocoel, 27 genes had higher expression levels and 45 genes had lower expression levels in infected mosquitoes compared to the uninfected control mosquitoes (Table S3). The gene CPIJ007162, which contains a predicted N-acetylmuramoyl-L-alanine amidase activity domain with innate immune function had four times lower expression levels in infected mosquitoes compared to the uninfected ones. Molecular functions enriched in genes with lower expression levels were related to carbamoyl-phosphate synthase and 4-alpha-hydroxytetrahydrobiopterin dehydratase activities (Figure S1). At this time point, GO terms associated with biological processes such as chloride transmembrane transport, chromatin organization and DNA packaging were enriched for genes with higher expression levels (Figure 3, Table S4).

### Transcriptional response at 10 days post blood feeding – sporozoite migration and invasion of salivary glands

At 10 dpf, after oocyst maturation and release of sporozoites as well as salivary glands invasion, 36 genes had lower expression levels and 37 genes had higher expression levels in infected mosquitoes (Table S5). An adhesive serine protease (CPIJ007535), a serine protease inhibitor (Serpin B8, CPIJ017784) and an uncharacterized gene (CPIJ009078) related to chitin metabolism had low expression levels at this time point. GO terms of molecular functions associated with ribonucleotide and GTP binding and oxidoreductase activity were enriched for genes with higher expression levels (Table S6, Figure S1). No GO terms were significantly enriched for genes with lower expression in infected mosquitoes.

## Discussion

We showed, for the first time, mosquito transcriptome responses to an avian malaria parasite. In this study, we tried to reproduce natural conditions as much as possible: 1) we used single wild-caught mosquitoes with minimal laboratory colonization (F1), 2) we obtained a “fresh” Hawaiian parasite isolate (only two passages in canaries), 3) and we used a donor bird with parasite intensity levels that *Culex* mosquitoes may be exposed to in Hawai□i (Woodworth et al., 2005). These attributes suggest that our results reflect real-world scenarios of *Culex*-*Plasmodium* interactions happening in Hawai□i. Indeed, serial parasite passages increase infection burden in birds and correlate with higher oocyst burden and longevity in mosquitoes (Pigeault et al., 2015), which are artificial effects that we aimed to avoid in our experiments. By simulating natural conditions while still benefiting from the controlled aspects of laboratory experiments, this approach allowed us to evaluate effects of *P. relictum* on Hawaiian *Cx. quinquefasciatus*. Not surprisingly, considering the profound physiological effects of blood-feeding on mosquitoes (Dana et al., 2005), we found that the time since exposure to blood had a larger overall effect on the mosquito transcriptome than the presence of *Plasmodium*. However, once those effects subsided, *Culex* transcriptional responses to *Plasmodium* infection were more pronounced during the early stages of parasite invasion than during development and sporozoite invasion of the salivary glands, which is similar to the generally strong immune response mounted by *Anopheles* mosquitoes during *Plasmodium* invasion of the midgut epithelia (Cirimotich et al., 2010).

Only two genes were differently expressed at more than one time point, suggesting that the *Culex* transcriptional response to *Plasmodium* is different across infection stages. Interestingly, a high proportion of *P. relictum* differently expressed genes are shared across different developmental stages in the vector (Sekar et al., 2021), revealing that mosquito transcriptional response to infection may not be coupled with the transcriptional changes in invading parasites.

Three genes involved in cytoskeleton organization had four times higher expression in infected mosquitoes at 24 hpf and no genes related to this biological process had lower expression rates at the same time point. Ookinete invasion of the midgut epithelia activates cytoskeleton reorganization in *Anopheles* (Vlachou et al., 2005), indicating this is a common feature of mosquito response to avian and mammalian *Plasmodium* during early stages of infection. Four genes with functions predicted to be involved in calcium transportation or binding had higher expression levels in infected mosquitoes at 24 hpf. Calcium is essential for ookinete motility (Bennink et al., 2016) and these alterations may supply parasites with Ca^2+^ which may facilitate midgut invasion. In *Anopheles* mosquitoes, increased nitric oxide synthase activity reduces parasite invasion after feeding on infected hosts (Luckhart et al., 1998), and gene transcription can increase as early as 6 hpf (Luckhart et al., 2003). In our study, oxidoreductase activity was enriched in infected mosquitoes at 24 hpf, encompassing 12 genes with lower expression rates in infected mosquitoes. However, we cannot determine whether this decreased activity would reduce the production of reactive nitrogen compounds that cause damage to *Plasmodium* parasites at early stages of infection (Luckhart et al., 1998). One of the three Down syndrome cell adhesion molecule gene paralogs had higher expression levels during ookinete invasion. This molecule mediates *Plasmodium* inhibition in *Anopheles* during ookinete migration through the midgut epithelia, before the formation of oocysts (Dong et al., 2012) and our results indicate that it may likewise participate in *Culex* response to avian malaria parasites.

At 24 hpf, we did not detect higher expression levels of genes related to apoptosis and cell immunity that are known to be overactivated during *Plasmodium* invasion in *Anopheles*, even though this stage of infection is marked by strong physiological and immune responses to infection (Rhodes & Michel, 2017). This may be because we used a host bird harboring a parasite intensity that translated to low numbers of parasites detected in the mosquitoes we dissected. Bird transcriptome response to avian malaria is highly dependent on blood parasite load (parasitemia), which may be related to pathogenicity (Videvall et al., 2020). These same authors found that low levels of circulating parasites were associated with a reduced transcriptome response to infection when compared with higher parasitemia. Further studies using vectors exposed to birds harboring different levels of parasitemia may clarify if mosquito response at the transcriptome level is proportional to the number of parasites undergoing development in the vector. Another reason we did not detect changes in the expression of these genes might be because the *Cx. quinquefasciatus* genome sequence is not as well assembled and annotated as those of *Anopheles gambiae* and *Aedes aegypti* (Giraldo-Calderón et al., 2015). In our study, 14.5% of *Cx. quinquefasciatus* DEGs in response to *P. relictum* infection were annotated as uncharacterized proteins or unspecified products with no predicted function, which limited our ability to interpret some of the results.

It has been observed that at the start and early acceleration of the sporogonic division, oocysts seem to be refractory to the mosquito immune response (Smith & Barillas-Mury, 2016), and this may explain the small transcriptome response to *Plasmodium* infection we observed at 5 dpf. However, some studies showed that mosquito immune response can destroy young oocysts because total numbers decrease during parasite development in *Anopheles* (Rhodes & Michel, 2017). The number of oocysts of avian *Plasmodium* also decrease during oocyst maturation in *Culex* (Sekar et al., 2021), and this may be due to immune-mediated parasite killing. An unspecified gene with a predicted peptidoglycan binding function (CPIJ007162) was downregulated at 5 dpf. Peptidoglycan recognition proteins (PGRP) are important in the regulation of the Immunodeficiency (Imd) pathway in insect midguts, a pathway shown to kill *Plasmodium* in *Anopheles* midguts (Mellroth et al., 2003). Some PGRP function as *P. falciparum* antagonists, reducing prevalence in *An. coluzzii* via Imd activation, while other PGRP genes from this family promote tolerance to *Plasmodium* by a downregulation of systemic Imd (Gendrin et al., 2017). This is the only DEG in our dataset with a predicted innate immune function, and we cannot infer whether the downregulation of this gene would affect *P. relictum* development within infected mosquitoes. Therefore, immune pathways in vector response to *Plasmodium* oocyst development remain to be described in *Culex* mosquitoes.

The mild transcriptome changes at 10 dpf indicate that a relatively small response is elicited in *Culex* to sporozoites during hemocoel migration and invasion of the salivary glands. This may be because salivary gland invasion by *Plasmodium* parasites generally induces lower mosquito physiological responses when compared to ookinete invasion of the midgut (Mueller et al., 2010). In *Anopheles* mosquitoes, some serine protease inhibitors, such as SRPN2, are known to facilitate *Plasmodium* midgut invasion and survival (Michel et al., 2005). SRPN6 is upregulated in the salivary glands of infected mosquitoes (Rosinski-Chupin et al., 2007) and reduce the number of invading sporozoites (Pinto et al., 2008). In contrast, we found that SRPN B8 (CPIJ017784) had lower expression levels in infected mosquitoes at 10 dpf, and future studies are warranted to investigate potential agonistic effects between this gene and *P. relictum* in *Culex*.

More than 80% of sporozoites released in the hemolymph are quickly destroyed before they invade the salivary glands (Hillyer et al., 2007). Drivers of mosquito response to circulating parasites are still unknown, and our experiment did not find specific immune-related genes that could be involved in sporozoite elimination. High nitric oxide (NO) concentrations driven by increased Nitric oxide synthase (NOS) expression reduce *Plasmodium* development at initial stages in the midgut (Luckhart et al., 2003). Expression levels of this gene are increased in the whole body of mosquitoes at the initiation of sporozoite release (Luckhart et al., 1998), but are not affected (Rosinski-Chupin et al., 2007) or can be reduced in the salivary glands during *Plasmodium* invasion (Dimopoulos et al., 1998). Here, a glutamate dehydrogenase gene, which has a predicted oxidoreductase function, had higher expression levels in infected *Culex* at 10 dpf, but we cannot infer if these changes translate into increased NO production in the body or salivary glands.

Although parasite invasion (24 hpf) was associated with a reduction in galactose catabolism and acetyl-CoA biosynthesis, parasite development (5 dpf and 10 dpf), did not elicit strong changes in the expression of gene groups involved in vector nutrient metabolism. We hypothesize that such changes are likely to be more evident in vectors with high parasite loads if disruption of mosquito nutrient metabolism is proportional to parasite burdens. Rodent malaria parasites trigger immune responses in *Anopheles* that reduce fitness and survival, while *P. falciparum* suppresses these responses, preventing the physiological cost of infection (reviewed by Shaw et al. 2021). Studies using a diversity of avian malaria parasites and bird-biting *Culex* mosquitoes found that *Plasmodium* infection does not reduce vector survival (Gutiérrez-López et al., 2020; Pigeault & Villa, 2018), or may presumably increase it (Gutiérrez-López et al., 2020; Vézilier et al., 2012) at the expense of reduced fecundity (Pigeault & Villa, 2018; Vézilier et al., 2012). In conclusion, the limited changes in the expression of genes related to nutrition metabolism and immune response we observed indicates that costs of infection for the vector may be minimal in the Hawaiian malaria system.

## Acknowledgements

We thank Eben Paxton, Elizabeth Abraham and many USGS crew members for the assistance in the field. This study was primarily funded by NSF Ecology and Evolution of Infectious Diseases Award 2001213 (PI DMF). Nicole Wagner was supported by the Rutgers NJ Agricultural Experiment Station and Elin Videvall was partly supported by a Swedish Research Council fellowship (2020-00259).

## Author contributions

Francisco C. Ferreira, Elin Videvall, Robert Fleischer and Dina Fonseca conceived and designed the study with inputs from all authors. Christa Seidl and Francisco C. Ferreira conducted field work and experimental infections. A. Marm Kilpatrick collected mosquito rafts in Hawai□i. Nicole E. Wagner and Francisco C. Ferreira conducted laboratory work. Elin Videvall and Francisco C. Ferreira curated and analyzed data. Dina Fonseca, Robert Fleischer and A. Marm Kilpatrick were responsible for funding acquisition and project administration. Francisco C. Ferreira led the writing of the manuscript with critical contributions from all authors.

## Data accessibility

Supporting information will be available online. Sequences have been uploaded to the Sequence Read Archive (SRA) at NCBI under accession number: PRJNA779986

## Figure legends

**Figure S1. Enriched Gene Ontology terms for molecular functions among differentially expressed genes in infected mosquitoes compared to uninfected ones**. GO terms for genes with higher expression rates in infected mosquitoes at 24 h post-feeding were almost significantly enriched (FDR = 0.107) and are displayed. No GO terms were enriched for genes with lower expression in infected mosquitoes at 10 days post feeding. hpf = hours post feeding, dpf = days post feeding. *oxidoreductase activity, acting on the CH-OH group of donors, NAD or NADP as acceptor; **oxidoreductase activity, acting on the CH-NH2 group of donors, NAD or NADP as acceptor.

## References

Aardema, M. L., Campana, M. G., Wagner, N. E., Ferreira, F. C., & Fonseca, D. M. (2021). A gene-based capture assay for surveying patterns of genetic diversity and insecticide resistance in a worldwide group of invasive mosquitoes. BioRxiv, 2021.08.24.457535. https://doi.org/10.1101/2021.08.24.457535

Anders, S., Pyl, P. T., & Huber, W. (2015). HTSeq—A Python framework to work with high-throughput sequencing data. Bioinformatics (Oxford, England), 31(2), 166–169. https://doi.org/10.1093/bioinformatics/btu638

Atkinson, C. T., & LaPointe, D. A. (2009). Introduced avian diseases, climate change, and the future of Hawaiian honeycreepers. Journal of Avian Medicine and Surgery, 23(1), 53–63. https://doi.org/10.1647/2008-059.1

Atkinson, C. T., Saili, K. S., Utzurrum, R. B., & Jarvi, S. I. (2013). Experimental evidence for evolved tolerance to avian malaria in a wild population of low elevation Hawai’i ‘Amakihi (Hemignathus virens). EcoHealth, 10(4), 366–375. https://doi.org/10.1007/s10393-013-0899-2

Bennink, S., Kiesow, M. J., & Pradel, G. (2016). The development of malaria parasites in the mosquito midgut. Cellular Microbiology, 18(7), 905–918. https://doi.org/10.1111/cmi.12604

Böhme, U., Otto, T. D., Cotton, J. A., Steinbiss, S., Sanders, M., Oyola, S. O., Nicot, A., Gandon, S., Patra, K. P., Herd, C., Bushell, E., Modrzynska, K. K., Billker, O., Vinetz, J. M., Rivero, A., Newbold, C. I., & Berriman, M. (2018). Complete avian malaria parasite genomes reveal features associated with lineage-specific evolution in birds and mammals. Genome Research, 28(4), 547–560. https://doi.org/10.1101/gr.218123.116

Bolger, A. M., Lohse, M., & Usadel, B. (2014). Trimmomatic: A flexible trimmer for Illumina sequence data. Bioinformatics, 30(15), 2114–2120. https://doi.org/10.1093/bioinformatics/btu170

Bonnot, T., Gillard, M. B., & Nagel, D. H. (2019). A simple protocol for Informative visualization of enriched Gene Ontology terms. Bio-Protocol, 9(22), e3429. https://doi.org/10.21769/BioProtoc.3429

Carlson, J. S., Giannitti, F., Valkiūnas, G., Tell, L. A., Snipes, J., Wright, S., & Cornel, A. J. (2016). A method to preserve low parasitaemia Plasmodium-infected avian blood for host and vector infectivity assays. Malaria Journal, 15, 154. https://doi.org/10.1186/s12936-016-1198-5

Cassin-Sackett, L., Callicrate, T. E., & Fleischer, R. C. (2019). Parallel evolution of gene classes, but not genes: Evidence from Hawai’ian honeycreeper populations exposed to avian malaria. Molecular Ecology, 28(3), 568–583. https://doi.org/10.1111/mec.14891

Cirimotich, C. M., Dong, Y., Garver, L. S., Sim, S., & Dimopoulos, G. (2010). Mosquito immune defenses against Plasmodium infection. Developmental and Comparative Immunology, 34(4), 387–395. https://doi.org/10.1016/j.dci.2009.12.005

Dana, A. N., Hong, Y. S., Kern, M. K., Hillenmeyer, M. E., Harker, B. W., Lobo, N. F., Hogan, J. R., Romans, P., & Collins, F. H. (2005). Gene expression patterns associated with blood-feeding in the malaria mosquito Anopheles gambiae. BMC Genomics, 6, 5. https://doi.org/10.1186/1471-2164-6-5

Dimopoulos, G., Seeley, D., Wolf, A., & Kafatos, F. C. (1998). Malaria infection of the mosquito Anopheles gambiae activates immune-responsive genes during critical transition stages of the parasite life cycle. The EMBO Journal, 17(21), 6115–6123. https://doi.org/10.1093/emboj/17.21.6115

Dine, D. L. V. (1904). Mosquitoes in Hawaii. In Bulletin of the Hawaii Agricultural Experimental Station (Vol. 6, p. 29).

Dong, Y., Cirimotich, C. M., Pike, A., Chandra, R., & Dimopoulos, G. (2012). Anopheles NF-κB-regulated splicing factors direct pathogen-specific repertoires of the hypervariable pattern recognition receptor AgDscam. Cell Host & Microbe, 12(4), 521–530. https://doi.org/10.1016/j.chom.2012.09.004

Ewels, P., Magnusson, M., Lundin, S., & Käller, M. (2016). MultiQC: Summarize analysis results for multiple tools and samples in a single report. Bioinformatics (Oxford, England), 32(19), 3047–3048. https://doi.org/10.1093/bioinformatics/btw354

Fecchio, A., Clark, N. J., Bell, J. A., Skeen, H. R., Lutz, H. L., De La Torre, G. M., Vaughan, J. A., Tkach, V. V., Schunck, F., Ferreira, F. C., Braga, É. M., Lugarini, C., Wamiti, W., Dispoto, J. H., Galen, S. C., Kirchgatter, K., Sagario, M. C., Cueto, V. R., González-Acuña, D., … Wells, K. (2021). Global drivers of avian haemosporidian infections vary across zoogeographical regions. Global Ecology and Biogeography, n/a(n/a). https://doi.org/10.1111/geb.13390

Feng, X., López Del Amo, V., Mameli, E., Lee, M., Bishop, A. L., Perrimon, N., & Gantz, V. M. (2021). Optimized CRISPR tools and site-directed transgenesis towards gene drive development in Culex quinquefasciatus mosquitoes. Nature Communications, 12(1), 2960. https://doi.org/10.1038/s41467-021-23239-0

Ferreira, F. C., Santiago-Alarcon, D., & Braga, É. M. (2020). Diptera Vectors of Avian Haemosporidians: With Emphasis on Tropical Regions. In D. Santiago-Alarcon & A. Marzal (Eds.), Avian Malaria and Related Parasites in the Tropics: Ecology, Evolution and Systematics (pp. 185–250). Springer International Publishing. https://doi.org/10.1007/978-3-030-51633-8_6

Fonseca, D. M., LaPointe, D. A., & Fleischer, R. C. (2000). Bottlenecks and multiple introductions: Population genetics of the vector of avian malaria in Hawaii. Molecular Ecology, 9(11), 1803–1814. https://doi.org/10.1046/j.1365-294x.2000.01070.x

Fonseca, D. M., Smith, J. L., Wilkerson, R. C., & Fleischer, R. C. (2006). Pathways of expansion and multiple introductions illustrated by large genetic differentiation among worldwide populations of the southern house mosquito. The American Journal of Tropical Medicine and Hygiene, 74(2), 284–289.

Gendrin, M., Turlure, F., Rodgers, F. H., Cohuet, A., Morlais, I., & Christophides, G. K. (2017). The Peptidoglycan Recognition Proteins PGRPLA and PGRPLB Regulate Anopheles Immunity to Bacteria and Affect Infection by Plasmodium. Journal of Innate Immunity, 9(4), 333–342. https://doi.org/10.1159/000452797

Giraldo-Calderón, G. I., Emrich, S. J., MacCallum, R. M., Maslen, G., Dialynas, E., Topalis, P., Ho, N., Gesing, S., VectorBase Consortium, Madey, G., Collins, F. H., & Lawson, D. (2015). VectorBase: An updated bioinformatics resource for invertebrate vectors and other organisms related with human diseases. Nucleic Acids Research, 43(Database issue), D707–713. https://doi.org/10.1093/nar/gku1117

Gutiérrez-López, R., Martínez-de la Puente, J., Gangoso, L., Soriguer, R., & Figuerola, J. (2020). Plasmodium transmission differs between mosquito species and parasite lineages. Parasitology, 147(4), 441–447. https://doi.org/10.1017/S0031182020000062

Hellgren, O., Waldenström, J., & Bensch, S. (2004). A New PCR assay for simultaneous studies of Leucocytozoon, Plasmodium, and Haemoproteus from avian blood. Journal of Parasitology, 90(4), 797–802. https://doi.org/10.1645/GE-184R1

Hillyer, J. F., Barreau, C., & Vernick, K. D. (2007). Efficiency of salivary gland invasion by malaria sporozoites is controlled by rapid sporozoite destruction in the mosquito hemocoel. International Journal for Parasitology, 37(6), 673–681. https://doi.org/10.1016/j.ijpara.2006.12.00

Huff, C. G. (1927). Studies on the Infectivity of Plasmodia of Birds for Mosquitoes, with special Reference to the Problem of Immunity in the Mosquito. American Journal of Hygiene, 7(6). https://www.cabdirect.org/cabdirect/abstract/19281000192

Kazlauskienė, R., Bernotienė, R., Palinauskas, V., Iezhova, T. A., & Valkiūnas, G. (2013). Plasmodium relictum (lineages pSGS1 and pGRW11): Complete synchronous sporogony in mosquitoes Culex pipiens pipiens. Experimental Parasitology, 133(4), 454–461. https://doi.org/10.1016/j.exppara.2013.01.008

Kim, D., Paggi, J. M., Park, C., Bennett, C., & Salzberg, S. L. (2019). Graph-based genome alignment and genotyping with HISAT2 and HISAT-genotype. Nature Biotechnology, 37(8), 907–915. https://doi.org/10.1038/s41587-019-0201-4

LaPointe, D. A., Goff, M. L., & Atkinson, C. T. (2005). Comparative susceptibility of introduced forest-dwelling mosquitoes in Hawai’i to avian malaria, Plasmodium relictum. Journal of Parasitology, 91(4), 843–849. https://doi.org/10.1645/GE-3431.1

LaPointe, D. A., Goff, M. L., & Atkinson, C. T. (2010). Thermal Constraints to the Sporogonic Development and Altitudinal Distribution of Avian Malaria Plasmodium relictum in Hawai’i. Journal of Parasitology, 96(2), 318–324. https://doi.org/10.1645/GE-2290.1

Li, H., Handsaker, B., Wysoker, A., Fennell, T., Ruan, J., Homer, N., Marth, G., Abecasis, G., Durbin, R., & 1000 Genome Project Data Processing Subgroup. (2009). The Sequence Alignment/Map format and SAMtools. Bioinformatics, 25(16), 2078–2079. https://doi.org/10.1093/bioinformatics/btp352

Love, M. I., Huber, W., & Anders, S. (2014). Moderated estimation of fold change and dispersion for RNA-seq data with DESeq2. Genome Biology, 15(12), 550. https://doi.org/10.1186/s13059-014-0550-8

Luckhart, S., Crampton, A. L., Zamora, R., Lieber, M. J., Dos Santos, P. C., Peterson, T. M. L., Emmith, N., Lim, J., Wink, D. A., & Vodovotz, Y. (2003). Mammalian transforming growth factor beta1 activated after ingestion by Anopheles stephensi modulates mosquito immunity. Infection and Immunity, 71(6), 3000–3009. https://doi.org/10.1128/IAI.71.6.3000-3009.2003

Luckhart, S., Vodovotz, Y., Cui, L., & Rosenberg, R. (1998). The mosquito Anopheles stephensi limits malaria parasite development with inducible synthesis of nitric oxide. Proceedings of the National Academy of Sciences of the United States of America, 95(10), 5700–5705. https://doi.org/10.1073/pnas.95.10.5700

McClure, K. M., Fleischer, R. C., & Kilpatrick, A. M. (2020). The role of native and introduced birds in transmission of avian malaria in Hawaii. Ecology, 101(7), e03038. https://doi.org/10.1002/ecy.3038

Mellroth, P., Karlsson, J., & Steiner, H. (2003). A scavenger function for a Drosophila peptidoglycan recognition protein. The Journal of Biological Chemistry, 278(9), 7059–7064. https://doi.org/10.1074/jbc.M208900200

Michel, K., Budd, A., Pinto, S., Gibson, T. J., & Kafatos, F. C. (2005). Anopheles gambiae SRPN2 facilitates midgut invasion by the malaria parasite Plasmodium berghei. EMBO Reports, 6(9), 891–897. https://doi.org/10.1038/sj.embor.7400478

Mueller, A.-K., Kohlhepp, F., Hammerschmidt, C., & Michel, K. (2010). Invasion of mosquito salivary glands by malaria parasites: Prerequisites and defense strategies. International Journal for Parasitology, 40(11), 1229–1235. https://doi.org/10.1016/j.ijpara.2010.05.005

Nishimoto, J. H. K. (2019). Integration of a “Self-docking Site” Genetic Construct in the Southern House Mosquito (Culex quinquefasciatus) as a Step Toward Genetic Control Strategies [M.S. Dissertation]. University of Hawai’i at Hilo.

Pacheco, M. A., Matta, N. E., Valkiunas, G., Parker, P. G., Mello, B., Stanley, C. E., Lentino, M., Garcia-Amado, M. A., Cranfield, M., Kosakovsky Pond, S. L., & Escalante, A. A. (2018). Mode and Rate of Evolution of Haemosporidian Mitochondrial Genomes: Timing the Radiation of Avian Parasites. Molecular Biology and Evolution, 35(2), 383–403. https://doi.org/10.1093/molbev/msx285

Paxton, E. H., Laut, M., Vetter, J. P., & Kendall, S. J. (2018). Research and management priorities for Hawaiian forest birds. The Condor, 120(3), 557–565. https://doi.org/10.1650/CONDOR-18-25.1

Pigeault, R., Vézilier, J., Cornet, S., Zélé, F., Nicot, A., Perret, P., Gandon, S., & Rivero, A. (2015). Avian malaria: A new lease of life for an old experimental model to study the evolutionary ecology of Plasmodium. Philosophical Transactions of the Royal Society B: Biological Sciences, 370(1675), 20140300. https://doi.org/10.1098/rstb.2014.0300

Pigeault, R., & Villa, M. (2018). Long-term pathogenic response to Plasmodium relictum infection in Culex pipiens mosquito. PLoS ONE, 13(2), e0192315. https://doi.org/10.1371/journal.pone.0192315

Pinto, S. B., Kafatos, F. C., & Michel, K. (2008). The parasite invasion marker SRPN6 reduces sporozoite numbers in salivary glands of Anopheles gambiae. Cellular Microbiology, 10(4), 891–898. https://doi.org/10.1111/j.1462-5822.2007.01091.x

R Development Core Team. (2020). R: A Language and Environment for Statistical Computing. R Foundation for Statistical Computing. http://www.R-project.org

Reidenbach, K. R., Cook, S., Bertone, M. A., Harbach, R. E., Wiegmann, B. M., & Besansky, N. J. (2009). Phylogenetic analysis and temporal diversification of mosquitoes (Diptera: Culicidae) based on nuclear genes and morphology. BMC Evolutionary Biology, 9, 298. https://doi.org/10.1186/1471-2148-9-298

Rhodes, V. L. M., & Michel, K. (2017). Chapter 4—Modulation of Mosquito Immune Defenses as a Control Strategy. In S. K. Wikel, S. Aksoy, & G. Dimopoulos (Eds.), Arthropod Vector: Controller of Disease Transmission, Volume 1 (pp. 59–89). Academic Press. https://doi.org/10.1016/B978-0-12-805350-8.00004-0

Rosinski-Chupin, I., Briolay, J., Brouilly, P., Perrot, S., Gomez, S. M., Chertemps, T., Roth, C. W., Keime, C., Gandrillon, O., Couble, P., & Brey, P. T. (2007). SAGE analysis of mosquito salivary gland transcriptomes during Plasmodium invasion. Cellular Microbiology, 9(3), 708–724. https://doi.org/10.1111/j.1462-5822.2006.00822.x

Sekar, V., Rivero, A., Pigeault, R., Gandon, S., Drews, A., Ahren, D., & Hellgren, O. (2021). Gene regulation of the avian malaria parasite Plasmodium relictum, during the different stages within the mosquito vector. Genomics, 113(4), 2327–2337. https://doi.org/10.1016/j.ygeno.2021.05.021

Shaw, W. R., Marcenac, P., & Catteruccia, F. (2021). Plasmodium development in Anopheles: A tale of shared resources. Trends in Parasitology, S1471-4922(21)00207-5. https://doi.org/10.1016/j.pt.2021.08.009

Smith, R. C., & Barillas-Mury, C. (2016). Plasmodium Oocysts: Overlooked Targets of Mosquito Immunity. Trends in Parasitology, 32(12), 979–990. https://doi.org/10.1016/j.pt.2016.08.012

Supek, F., Bošnjak, M., Škunca, N., & Šmuc, T. (2011). REVIGO summarizes and visualizes long lists of gene ontology terms. PloS One, 6(7), e21800. https://doi.org/10.1371/journal.pone.0021800

Valkiūnas, G. (2005). Avian Malaria Parasites and Other Haemosporidia (1st edition). CRC Press.

Valkiūnas, G., & Iezhova, T. A. (2017). Exo-erythrocytic development of avian malaria and related haemosporidian parasites. Malaria Journal, 16. https://doi.org/10.1186/s12936-017-1746-7

van Riper, Charles. I., van Riper, S. G., Goff, M. L., & Laird, M. (1986). The Epizootiology and Ecological Significance of Malaria in Hawaiian Land Birds. Ecological Monographs, 56(4), 327–344. https://doi.org/10.2307/1942550

Vézilier, J., Nicot, A., Gandon, S., & Rivero, A. (2012). Plasmodium infection decreases fecundity and increases survival of mosquitoes. Proceedings. Biological Sciences, 279(1744), 4033–4041. https://doi.org/10.1098/rspb.2012.1394

Videvall, E., Palinauskas, V., Valkiūnas, G., & Hellgren, O. (2020). Host Transcriptional Responses to High-and Low-Virulent Avian Malaria Parasites. The American Naturalist, 195(6), 1070–1084. https://doi.org/10.1086/708530

Videvall, E., Paxton, K. L., Campana, M. G., Cassin-Sackett, L., Atkinson, C. T., & Fleischer, R. C. (2021). Transcriptome assembly and differential gene expression of the invasive avian malaria parasite Plasmodium relictum in Hawai’i. Ecology and Evolution, 11(9), 4935–4944. https://doi.org/10.1002/ece3.7401

Vlachou, D., Schlegelmilch, T., Christophides, G. K., & Kafatos, F. C. (2005). Functional Genomic Analysis of Midgut Epithelial Responses in Anopheles during Plasmodium Invasion. Current Biology, 15(13), 1185–1195. https://doi.org/10.1016/j.cub.2005.06.044

Warner, R. E. (1968). The role of introduced diseases in the extinction of the endemic Hawaiian avifauna. The Condor, 70(2), 101–120.

Woodworth, B. L., Atkinson, C. T., Lapointe, D. A., Hart, P. J., Spiegel, C. S., Tweed, E. J., Henneman, C., Lebrun, J., Denette, T., Demots, R., Kozar, K. L., Triglia, D., Lease, D., Gregor, A., Smith, T., & Duffy, D. (2005). Host population persistence in the face of introduced vector-borne diseases: Hawaii amakihi and avian malaria. Proceedings of the National Academy of Sciences of the United States of America, 102(5), 1531–1536. https://doi.org/10.1073/pnas.0409454102

