## Supplementary figures and images for "Transcriptional response of individual Hawaiian *Culex quinquefasciatus* mosquitoes to the avian malaria parasite *Plasmodium relictum*"

### Figure S1

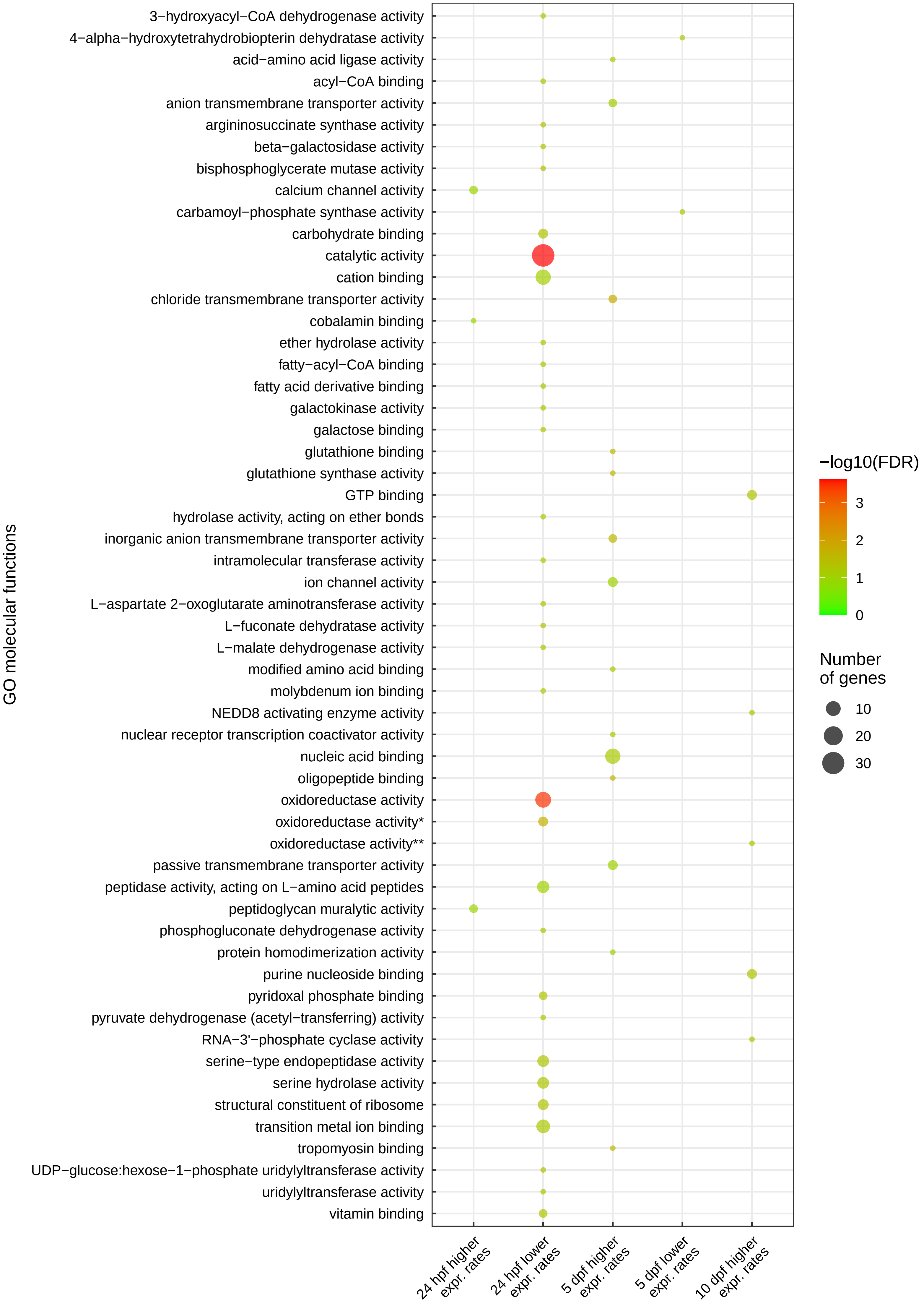
